# Ahead of Maturation: Enhanced Speech Envelope Training Boosts Rise Time Discrimination in Pre-Readers at Cognitive Risk for Dyslexia

**DOI:** 10.1101/2021.04.12.439411

**Authors:** Shauni Van Herck, Femke Vanden Bempt, Maria Economou, Jolijn Vanderauwera, Toivo Glatz, Benjamin Dieudonné, Maaike Vandermosten, Pol Ghesquière, Jan Wouters

**Author notes:** **Author note** Shauni Van Herck.

## Abstract

Dyslexia has frequently been related to atypical auditory temporal processing and speech perception. Results of studies emphasizing speech onset cues and reinforcing the temporal structure of the speech envelope, i.e. envelope enhancement, demonstrated reduced speech perception deficits in individuals with dyslexia. The use of this strategy as an auditory intervention might thus reduce some of the deficits related to dyslexia. Importantly, interventions are most effective when they are provided during kindergarten and first grade. Hence, we provided a tablet-based 12-week preventive auditory and phonics-based intervention to pre-readers at cognitive risk for dyslexia and investigated the effect on auditory temporal processing with a rise time discrimination task. Ninety-one pre-readers at cognitive risk for dyslexia (aged 5-6) were assigned to two groups receiving a phonics-based intervention and playing a story listening game either with (*n* = 31) or without (*n* = 31) envelope enhancement or a third group playing control games and listening to non-enhanced stories (*n* = 29). Rise time discrimination was measured directly before, directly after and one year after the intervention. While the groups listening to non-enhanced stories mainly improved after the intervention during first grade, the group listening to enhanced stories improved during the intervention in kindergarten and subsequently remained stable during first grade. Hence, an envelope enhancement intervention improves auditory processing skills important for the development of phonological skills. This occurred before the onset of reading instruction, preceding the maturational improvement of these skills, hence giving at risk children a head start when learning to read.

**Research highlights:**

- The first investigation of speech envelope enhancement as a potential preventive intervention strategy in pre-readers at cognitive risk for dyslexia
- Speech envelope enhancement increases the rise time sensitivity of children at cognitive risk for dyslexia
- Rise time discrimination can be enhanced before formal reading instruction, a crucial period in development

---

Developmental dyslexia is a learning disability, which manifests through severe and persistent difficulties in reading and/or spelling which cannot be explained by low intelligence, insufficient instruction or sensory disabilities (Peterson & Pennington, 2012; Vellutino et al., 2004). Dyslexia has a multifactorial etiology with several cognitive and neural factors contributing to its development (Menghini et al., 2010; Pennington, 2006; van Bergen et al., 2014). Within this multifactorial framework, deviant phonological processing has been considered a core deficit (Ramus et al., 2003) and therefore phonics instruction is considered the best approach for learning to read (National Institute of Child Health and Human Development, 2000). By some, this phonological deficit has been assigned to difficulties with auditory perception, allocated to the processing of temporal information (i.e. changes over time) (Farmer & Klein, 1995; Goswami, 2011). Initial theories suggesting auditory deficits focused primarily on rapid auditory processing deficits caused by difficulties with processing brief auditory stimuli or auditory stimuli presented in rapid succession (Farmer & Klein, 1995; Tallal, 1980). More recently, this focus has shifted to a deficit in processing the slow-rate dynamic information in speech (Goswami et al., 2002; Poelmans et al., 2011; Richardson et al., 2004). The latter studies emphasize the importance of amplitude rise times in the speech envelope. The speech envelope contains modulations in the speech signal between 2 and 50 Hz (Rosen, 1992). Amplitude rise times indicate the time it takes to reach peak amplitude of an amplitude modulation in the speech envelope (Goswami, 2019). A cascading effect can explain how temporal cues such as rise times contribute to (the development of) dyslexia. Rise times provide cues for the segmentation of the speech signal into its constituting elements such as syllables and phonemes. This process is mediated by entrainment to the speech signal, for which rise times provide important cues considering their information on the different rates of amplitude modulations in the speech signal (Goswami, 2011; Gross et al., 2013; Peelle & Davis, 2012). This entrainment is known to aid speech segmentation and intelligibility (Doelling et al., 2014; Giraud & Poeppel, 2012; Gross et al., 2013; Luo & Poeppel, 2007), but it has been suggested to be compromised in dyslexia (Goswami, 2011). As such, the failure to accurately and efficiently use rise time cues by people with dyslexia would impact neural entrainment, speech perception, and consequently phonological and reading development (Goswami et al., 2010; Power et al., 2016; Ziegler et al., 2009). At the behavioral level, difficulties with discriminating rise times have been established in both adults (Hämäläinen et al., 2005; Law et al., 2014; Leong et al., 2011) and children with dyslexia (Goswami et al., 2002; Goswami et al., 2011; Law, Wouters, et al., 2017; Poelmans et al., 2011; Richardson et al., 2004; Vanvooren et al., 2017). Even preceding the start of formal reading instruction rise time discrimination (RTD) can differentiate between children at high family risk for dyslexia and control children at low family risk for dyslexia (Law, Wouters, et al., 2017). Furthermore, a study by Law et al. (2014) revealed rise time discrimination deficits in 58% of the dyslexic adult participants in their study, highlighting the importance of auditory processing in dyslexia. At the neurophysiological level, a study by Van Hirtum, Ghesquière, & Wouters (2019) also revealed an impaired tracking of rise time cues in young adults with dyslexia by comparing neural synchronization to stimuli with differing rise times. Apart from deficits in rise time discrimination at both the behavioral and neurophysiological level, clear links have been established between rise time discrimination and reading-related skills such as phonological awareness, even preceding the onset of formal reading instruction (Goswami et al., 2020; Vanvooren et al., 2017). A longitudinal study by Vanvooren et al. (2017) established a link between pre-reading children’s performance on rise time discrimination and later phonology and literacy performance. This study confirmed that basic auditory processing already exerts an influence on phonology and literacy before the onset of formal reading instruction and that these basic auditory processes like rise time discrimination have a profound impact on reading development. Goswami et al. (2020) even postulated rise time discrimination to be the core contributor to the development of phonological awareness. For rise time discrimination to affect phonological awareness through speech perception abilities, there should be a link established between basic auditory processing such as the processing of rise time cues and speech perception. Even though this link has not always been clearly demonstrated (Law et al., 2014; Poelmans et al., 2011), two recent studies by Van Hirtum, Moncada-Torres, et al. (2019) and Van Hirtum et al. (in press) revealed that difficulties with processing temporal cues in the speech envelope have a considerable impact on the speech perception deficit in both children and adults with dyslexia. The researchers emphasized rise times in the speech envelope to facilitate their detection, and investigated the impact on speech perception in a speech in noise task with vocoded speech, in which there is only limited temporal fine structure and spectral information, and in natural speech. Enhancing the rise times was found to normalize speech perception in this task. This effect occurred passively (i.e. without explicit instruction) and instantaneously (i.e. after only limited exposure) and results were similar in the natural and vocoded speech conditions. Hence, speech perception deficits in dyslexia indeed seem to stem from processing envelope cues. The method used in the latter studies in which rise times were amplified is referred to as Envelope Enhancement (EE), originally developed for use in cochlear implant users (Geurts & Wouters, 1999; Koning & Wouters, 2012, 2016). Emphasizing rise time cues in the speech envelope allegedly enhances the suboptimal entrainment of neural oscillations to the speech signal in people with dyslexia, thereby improving their speech perception (Doelling et al., 2014; Gross et al., 2013; Van Hirtum, Moncada-Torres, et al., 2019; Van Hirtum et al., in press). Considering its efficiency in children with dyslexia using a speech in noise task and the importance of speech perception for phonological and reading development, EE has been suggested as a potential intervention strategy (Van Hirtum et al., in press).

A meta-analysis by Wanzek & Vaughn (2007) revealed the highest effects for early interventions administered during kindergarten and first grade, so interventions should preferably be provided at an early age. Yet, dyslexia is usually only diagnosed after several years of reading/writing instruction, when reading and writing impairments show to be severe (usually in second grade or later) and hence interventions are mainly provided after the period in which they would be most effective. This paradox is called the ‘dyslexia paradox’ (Ozernov-Palchik & Gaab, 2016) and clearly indicates the importance of preventive intervention. Meta-analyses have provided most support for phonics-based interventions, especially when they include explicit and systematic phonics instruction and when they are provided at an early age (Bus & Van Ijzendoorn, 1999; National Institute of Child Health and Human Development, 2000; Richardson & Lyytinen, 2014; Snowling & Hulme, 2011; Verhoeven et al., 2020). Despite the positive impact of these phonics-based interventions, children with dyslexia typically do not reach the reading level of their typical reading peers, even when the interventions are provided at an early age (McTigue et al., 2019). A possible explanation for this is that phonics-based interventions rely on the accurate perception of the presented phonemes. However, due to the auditory processing deficit in some people with dyslexia, children with dyslexia presumably have difficulties with this accurate perception of phonemes (Ramus et al., 2003; Vandermosten et al., 2010, 2011), and hence, might not sufficiently benefit from these interventions as to reach the reading level of typical readers. Auditory intervention strategies might provide a solution to this. Musical and rhythm-based interventions that train temporal processing and rhythm skills have already been shown to provide benefits to phonological and literacy development (Bhide et al., 2013; Degé & Schwarzer, 2011; Flaugnacco et al., 2015; Harrison et al., 2018). Apart from these auditory interventions, we believe that EE also shows potential as an intervention strategy. EE might enable children with dyslexia to benefit more from phonics-based interventions by normalizing speech perception and hence improving the perception of phonemes and consequently the phoneme representations. EE has proven to be effective during a speech in noise task in children with dyslexia, and does not require children to have any reading experience, making it particularly well suited for early intervention (Van Hirtum et al., in press).

Notwithstanding the promising results using EE during a speech in noise task in adults and children with dyslexia (Van Hirtum, Moncada-Torres et al., 2019; Van Hirtum et al., in press), its potential to be used as (preventive) intervention strategy has not yet been investigated. Considering its suggested impact on the efficacy of phonics-based interventions in people with auditory processing deficits, it might be an additional benefit to combine EE with a phonics-based intervention. Therefore, the goal of this study was to explore the additive auditory perceptive effects of a preventive intervention combining EE and phonics-based training in Dutch-speaking pre-readers at cognitive risk for dyslexia. Our 12-week preventive intervention combined a tablet-based Flemish version of GraphoGame, a frequently implemented phonics-based intervention initially developed for children from first grade struggling with acquiring basic decoding skills (McTigue et al., 2019; Richardson & Lyytinen, 2014), with an auditory story-game intervention in which the stories were envelope enhanced. We implemented an EE strategy adapted from the one of Koning & Wouters (2012). The outcome measure of this study is rise time discrimination, which has been proposed as a contributor to one of the core deficits in dyslexia, namely the sensitivity to temporal cues in the speech envelope (Farmer & Klein, 1995; Goswami, 2011; Goswami et al., 2002; Goswami et al., 2020; Hämäläinen et al., 2013; Poelmans et al., 2011; Van Hirtum, Ghesquière, & Wouters, 2019; Vanvooren et al., 2017). Considering the speech perception improvement by EE in both children and adults with dyslexia (Van Hirtum, Moncada-Torres, et al., 2019; Van Hirtum et al., in press), we expected EE to also boost basic auditory processing. Precisely, since EE specifically enhances rise times in the speech envelope, we hypothesize children at cognitive risk for dyslexia receiving an extensive EE training to exhibit enhanced rise time sensitivity directly after the intervention, with potential long term maintenance, whereas we do not expect GraphoGame to impact performance. To the best of our knowledge, no study has yet investigated the ability of EE to improve basic auditory processing in children at risk for dyslexia, prior to reading instruction. This could result in the development of more appropriate interventions targeting an important deficit in dyslexia.

## Materials and Methods

### Participants

Participants for the current study were recruited during a large-scale screening phase in children attending the last year of kindergarten. Parents of children attending kindergarten in Flanders (Belgium) were contacted by sending an information letter and video. From the 1900 children who received parental consent for participation in the screening, 1225 children were eventually assessed based on a few exclusion criteria. All children had to be native Dutch speakers with bilateral normal hearing, neither history of brain damage or neurological disorders, nor (preliminary) diagnosis of ASS or ADHD, and did not receive any therapy due to language and/or articulatory problems. Children with incomplete or unreliable test scores, as well as children scoring below the 10^th^ percentile on non-verbal IQ, i.e. below a norm score of 75.3 on the Raven’s Colored Progressive Matrices (Raven et al., 1984), were further excluded, leaving us with a sample of 1091 children. From these 1091 children, 119 children were eventually defined as having an elevated cognitive risk for dyslexia, defined as scoring below the 30^th^ percentile on two out of three reading-related measures. These measures were phonological awareness (PA), letter knowledge (LK) and rapid automatized naming (RAN), the strongest cognitive pre-literacy predictors of dyslexia (Caravolas et al., 2012; Clayton et al., 2020; Ozernov-Palchik & Gaab, 2016). Children scoring below the 30^th^ percentile on PA and RAN were additionally required to have LK below the 40^th^ percentile. A detailed description of the screening tasks and procedure can be found in Verwimp et al. (2020). From the 119 children at cognitive risk for developing dyslexia, 91 participated in the current study. Eight children were lost to post-test and 12 children were lost to consolidation test, resulting in a sample of 83 and 79 children at post-test and consolidation test respectively. Seventy-six of the children have all three test phases complete. However, data from all 91 children are included for one or more test phases, since statistical analyses are robust to missing data. Due to technical issues with the tablets used for the intervention, EE data from 4 children were lost. The study was approved by the ethics committee of the University Hospitals of Leuven and written parental consent as well as oral assent by the children was obtained for all the subjects before participation.

### Procedure

The 91 selected children participated in a longitudinal study with a pre-test, intervention, post-test, consolidation test design. They performed an initial baseline measurement of the outcome measure (i.e. rise time discrimination). Afterwards, the children were instructed to carry out the tablet-based intervention at home in which the children played tablet games for six days a week for a total period of 12 weeks. For the EE intervention children listened to one to two stories per day, resulting in approximately 10 minutes of story-listening, while GraphoGame had to be played for 15 minutes per day. Not all children played the games according to our instructions (i.e. 6 days per week for 12 weeks), so intervention duration was not strictly 12 weeks for all children, but rather on average 14 weeks. For the EE intervention children completed on average 91% of the intervention, while for GraphoGame on average 89% was completed. To complete the intervention, all children received a Samsung Galaxy Tab E9.6 tablet in combination with an Audiotechnica ATHM20x headphone. Our research group calibrated the tablets and headphones for sound level for the EE intervention. Following the intervention, children returned for a post-test in which they were again tested for rise time discrimination. Post-test took place on average 1.5 week after the end of the intervention. On average 1 year and 2 months after the post-test, after roughly one year of formal reading instruction in first grade, the children returned for a consolidation test in which their rise time discrimination was again investigated. Through a block randomization all children were randomly assigned to one of three experimental groups. All children were instructed to play the games for their respective groups for 6 days a week for a total period of 12 weeks. The first experimental group was instructed to perform a phonics-based training (i.e. GraphoGame, GG) for 15 minutes in combination with actively listening to stories that were envelope enhanced (EE) for 10 minutes (i.e. GGEE group; *n*_pre-test_ = 31, *n*_post-test_ = 29, *n*_consolidation test_ = 26). Children in the second experimental group also played GraphoGame for 15 minutes, but listened to stories without envelope enhancement (NE) for 10 minutes (i.e. GGNE group; *n*_pre-test_ = 31, *n*_post-test_ = 28, *n*_consolidation test_ = 29). Children in the third experimental group constituted the active control group. They played control games for 15 minutes and actively listened to stories without envelope enhancement (NE) for 10 minutes (i.e. ACNE group; *n*_pre-test_ = 29, *n*_post-test_ = 26, *n*_consolidation test_ = 24). A chi-squared test revealed no differences between the groups in terms of drop-out (χ^2^(2, *N*=12) = 1.5, *p* = .47). The block randomization assured that the three experimental groups were balanced with respect to birth trimester, sex, and educational environment (i.e. school) and did not differ in age (months), non-verbal IQ, sex and socio-economic status (SES) (see Table 1 for participant characteristics).

**Table 1.**
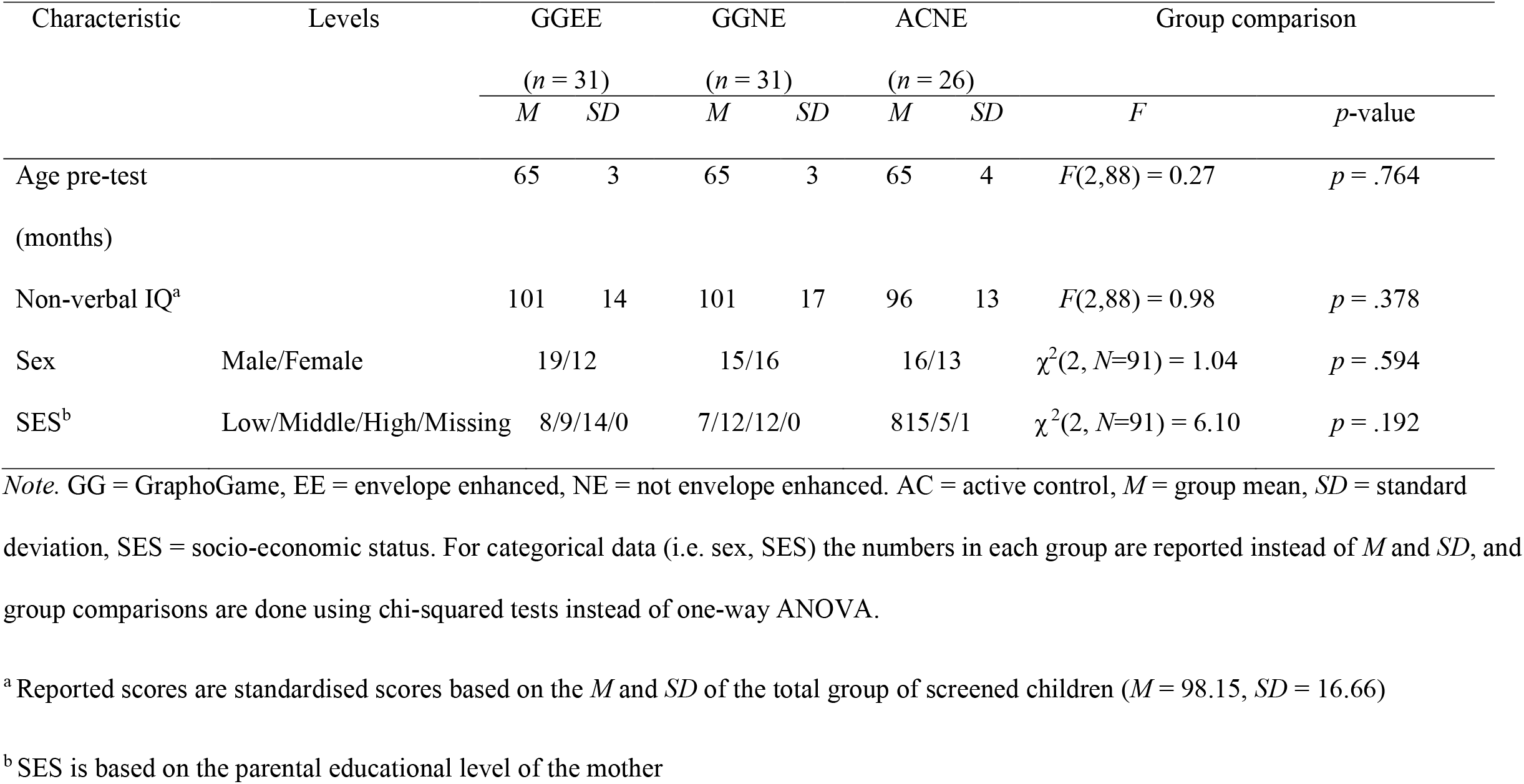
Participant characteristics.

### Interventions

#### Envelope Enhancement

Signal processing of 72 spoken stories was performed in MATLAB R2016b (The MathWorks Inc 2016). The signal processing methods here were adapted from those in the study of Koning & Wouters (2012). As a first step in the signal processing, the original signal *s*(*t*) was resampled to *f*_*sampling*_ = 16 kHz and subsequently split into frames of 128 samples with a frame advance of 32 samples. A frame-based fast Fourier transform preceded by Hann windowing transformed the signal to the time-frequency domain. Upon this, the frequency bins were combined by means of a weighted sum of their powers, mapping them to 23 critical channels. The frequency limits of these channels were defined according to the critical bandwidths of Fastl and Zwicker (2007). The envelope of each channel *E*(*t, k*) was obtained by taking the square root per channel. A slow envelope of each channel *E*_*slow*_(*t, k*) was obtained as follows. The envelope *E*(*t, k*) was low-pass filtered with a fourth-order Butterworth filter with a cutoff frequency of 20 Hz, which gave a signal with higher time delay when reacting to sudden increases in *E*(*t, k*). This low-pass filtered version of *E*(*t, k*) was then half-wave rectified and amplified by a factor of *A*_*slow*_ = 8 to obtain *E*_*slow*_(*t, k*). The latter step ensured a higher level of *E*_*slow*_(*t, k*) than that of *E*(*t, k*) at quasi-stationary parts, while the level of *E*(*t, k*) was higher than that of *E*_*slow*(*t,k*)_ at sudden increases in energy (i.e. onsets). As such, subtracting *E*_*slow*_(*t, k*) from *E*(*t, k*) yields a signal with peaks at the onsets of the envelope *E*(*t, k*), and negative values at stationary parts of *E*(*t, k*). A peak envelope signal *E*_*peak*_(*t, k*) could therefore be obtained by subtracting *E*_*slow*_(*t, k*) from *E*(*t, k*), followed by half-wave rectification and amplification by a factor of *A*_*peak*_ = 3.5. The eventual peak signal in the time domain *p*(*t*) was obtained by mapping the 23 bands back to 128 frequency bins, followed by an inverse fast Fourier transform after which the frames were recombined by a weighted overlap-add method. Finally, *p*(*t*) was added to the original signal *s*(*t*), resulting in the envelope enhanced signal *s*_*EE*_(*t*), see Figure 1.

**Figure 1.**
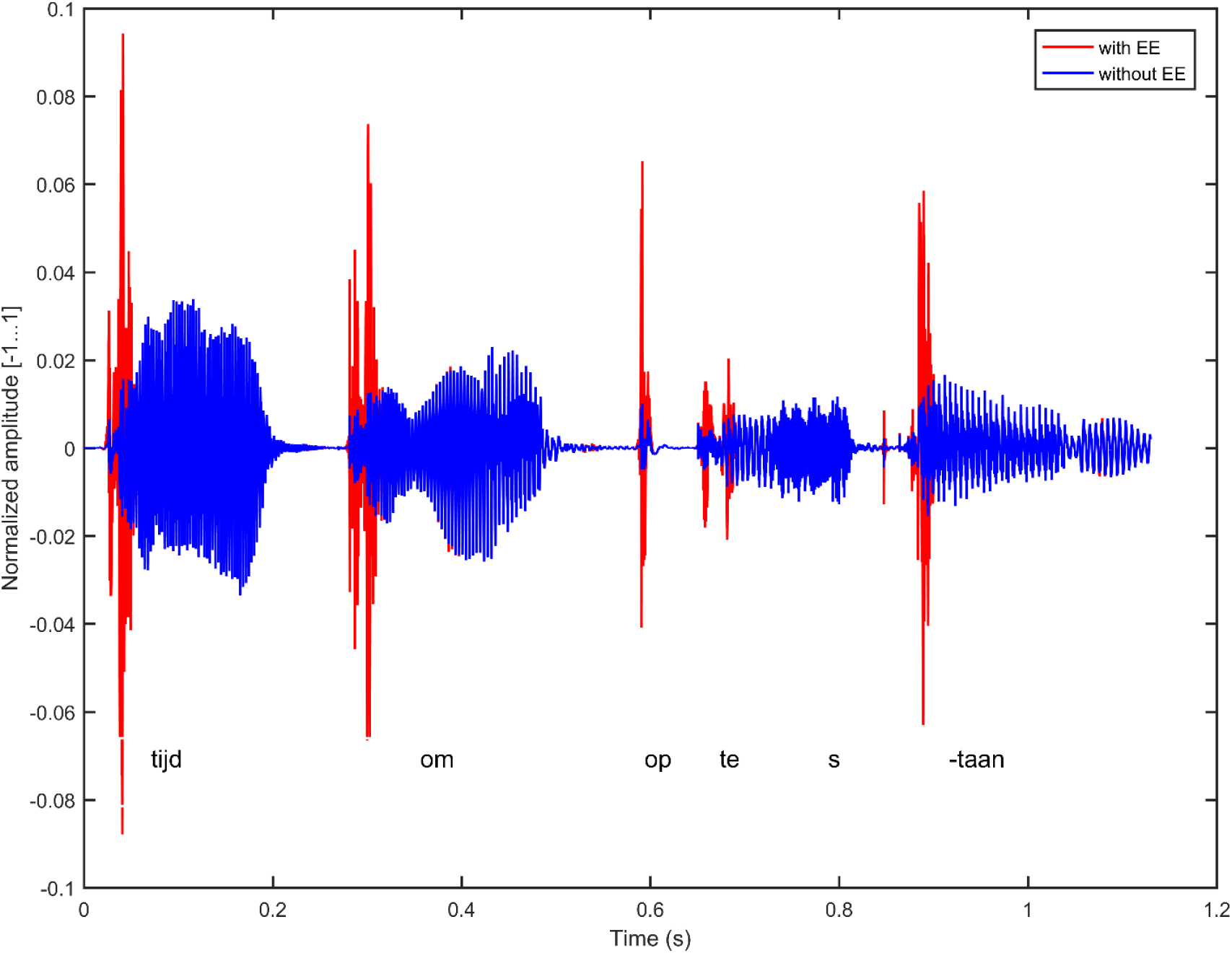
EE/NE Stimulus. *Note*. A waveform of a sentence from one of the stories without EE (blue) and with EE (red)

Eighty-seven age-appropriate stories ranging from 3 to 16 minutes were selected from 18 book series. Nine professionals with knowledge of standard Dutch recorded these stories. The stories were then combined with appropriate images and included in a tablet-game. Two versions of the game were created, one in which the stories were enhanced with the envelope enhancement signal processing strategy (EE) and one without enhancement (NE). The game consisted of 18 themes or book series that each contained 4 story sessions. Each story session contained one long or two short stories. The children played one story session per day. Hence, in total there were 72 playing days, covering the entire 12-week intervention period with one day off every week. At the end of each story, the children answered questions about its content to ensure active listening. To ensure continued motivation, children earned coins for every correct response, allowing them to buy gadgets for their game character. The children’s progress was logged online to allow the researchers to monitor it.

#### GraphoGame

GraphoGame Flemish (Glatz, 2021) consisted of mini-games divided over three module types namely, grapheme modules, phonology modules, and reading modules. The different mini-games aimed at training or assessing different tasks, such as introducing new graphemes, visual discrimination, auditory discrimination, phoneme blending, spelling, phoneme counting, reading, general purpose levels and playing motivation assessment. The game’s content was slowly increasing in difficulty, and was enough to span the extended 12-week intervention period. During the 12-week intervention, the children needed to progress through the game content, but they were able to go back to play previous levels at all times. To increase motivation, children earned coins when progressing though the game, which enabled them to buy gadgets for their game character in an in-game store. The results from the GraphoGame sessions were logged online and were accessible to the research group in order to follow up the children’s progress throughout the game. GraphoGame Flemish is described in detail in Glatz (2021).

#### Control Game

The control game consisted of six commercially available applications, namely Lego City My City, LegoDuploTown, LegoDuploTrains, Playmobile horseriding game, Playmobil Police and Lego Heartlake Rush. The children were not instructed to play any particular game, as long as they played according to the schedule (i.e. 6 days per week for a period of 12 weeks). Identical to GraphoGame, children played the control games for 15 minutes per day. The choice for these games was based on a few criteria. They (1) were at least as immersive as GraphoGame, (2) were age-appropriate, (3) were shown by an internal pilot study to not train reading, and to induce the same placebo-effect for attention span as GraphoGame.

### Measures: Pre-test, Post-test and Consolidation Test

#### Rise Time Discrimination Task

Auditory perception was investigated using a rise time discrimination task identical to the one used in Vanvooren et al. (2017), which measures the participants’ sensitivity to differences in rise times. The task was performed in a soundproof booth. Prior to performing the rise time discrimination task, all participants were screened for hearing loss by means of an audiometric pure-tone detection task at .25, .5, 1, 2, 4 and 8 kHz. The Fletcher Index (FI) was calculated as the average threshold at .5, 1 and 2 kHz (Fletcher, 1950). Stimuli were presented monaurally to the right ear over calibrated Sennheiser HDA 200 headphones at 70 dB SPL, unless the FI to the right ear exceeded 20. In this case, the stimuli were presented at 70 dB SPL + (FI-20 dB SPL). This was the case for three out of 91 children at pre-test, seven out of 83 at post-test, and six out of 79 at consolidation test. Rise time discrimination was estimated with a two-down, one-up adaptive staircase procedure, targeting the threshold corresponding to 70.7 % correct responses (Levitt, 1971). This procedure was embedded within a three-alternative forced-choice oddity paradigm and presented as an interactive computer gamification using APEX software (Francart et al., 2008; Laneau et al., 2005). A threshold run was terminated after eight reversals and the average of the last four reversals was taken as the threshold. Each participant completed the task two times, thus obtaining two thresholds. To guarantee understanding of the task, the participants completed a practice run prior to the two actual threshold runs. Hence, the participants were already familiarized with the task. Stimuli comprised speech-weighted noise with a linear rise and fall time. The total duration of the stimuli was fixed at 800 ms, including a fixed linear fall time of 75 ms. The rise times varied logarithmically between 15 and 699 ms in 50 steps, with the stimulus with the shortest rise time (15 ms) as the reference stimulus in each trial. The recorded threshold is the rise time of the target stimulus at which the children can still reliably discriminate between the target and the reference stimulus. Analyses were performed on the best threshold (i.e. the lowest threshold) of the two obtained thresholds (Law, Vandermosten, et al., 2017).

### Statistical analysis

All statistical analyses were performed in R (version 3.5.1) (R Core Team, 2018). Prior to analyses, the assumption of normality of the residuals was assessed using Kolmogorov-Smirnov tests. To ensure the assumption of homogeneity of variance, Levene tests were performed. For the rise time discrimination task, both the assumptions of normality of the residuals and homogeneity of variance were violated, steering us towards a semi-parametric approach for data analysis. This main analysis investigated the short- and long-term effects of the auditory and phonics-based interventions using generalized estimating equations (GEE) (geepack package) (Halekoh et al., 2006; Zeger & Liang, 1986). Based on the distribution of the dependent variable (i.e. RTD threshold), a Gamma Family with a log link, which exponentiates the linear predictors to make the model fit in a linear form, was specified. Considering the repeated measures design, an autoregressive correlation structure was specified, accounting for the correlation between repeated observations from a given subject. Missing values were reported to the model. For analyses, the interaction between intervention group and test phase was considered. In order to disentangle short- and long-term intervention effects between the different intervention groups, we performed post-hoc comparisons using estimated marginal means (EMMs) with a Holm correction for multiple comparisons. In an additional analysis, the number of hours the groups spent listening to the stories and the number of hours the groups played either GG or CG were investigated using One-way ANOVAs, since residuals were normally distributed for these measures. Considering the non-normality of the residuals for the rise time discrimination task, all descriptives are reported as medians and interquartile ranges, whereas intervention durations are reported as means and standard deviations.

## Results

Descriptive data for the RTD thresholds are provided in Table 2 and an illustration of the thresholds for all groups and test phases can be found in Figure 2. We observed a significant interaction effect between group and test phase (χ^2^(4) = 11, *p* = .026). The interaction effect showed that whereas all groups improved significantly from pre-test to consolidation test (GGEE: *z* = 6.05, *p* < .001, 95 % asymptotic CI [1.57, 4.33], GGNE: *z* = 4.72, *p* < .001, 95 % asymptotic CI [1.29, 3.75]; ACNE: *z* = 4.00, *p* = .002, 95 % asymptotic CI [1.22, 5.98]), only the GGEE group’s performance was significantly enhanced at post-test (*z* = 5.15, *p* < .001, 95 % asymptotic CI [1.39, 4.07]). An improvement in the GGNE group took place during 1^st^ grade (*z* = 3.12, *p* = 0.044, 95 % asymptotic CI [0.99, 2.89]), and the ACNE group’s improvement during this same time period came close to significance (*z* = 2.99, *p* = .063, 95 % asymptotic CI [0.95, 4.44]). Despite the differences between the groups regarding the period in which the main improvement took place, none of the groups differed from the other at the three test phases, although at post-test the difference between GGEE and ACNE almost reached significance with the ACNE group performing worse than the GGEE group (*z* = -2.94, *p* = .073, 95% asymptotic CI [0.28, 1.06]). See Figure 3 for the GEE model predictions.

**Table 2.**
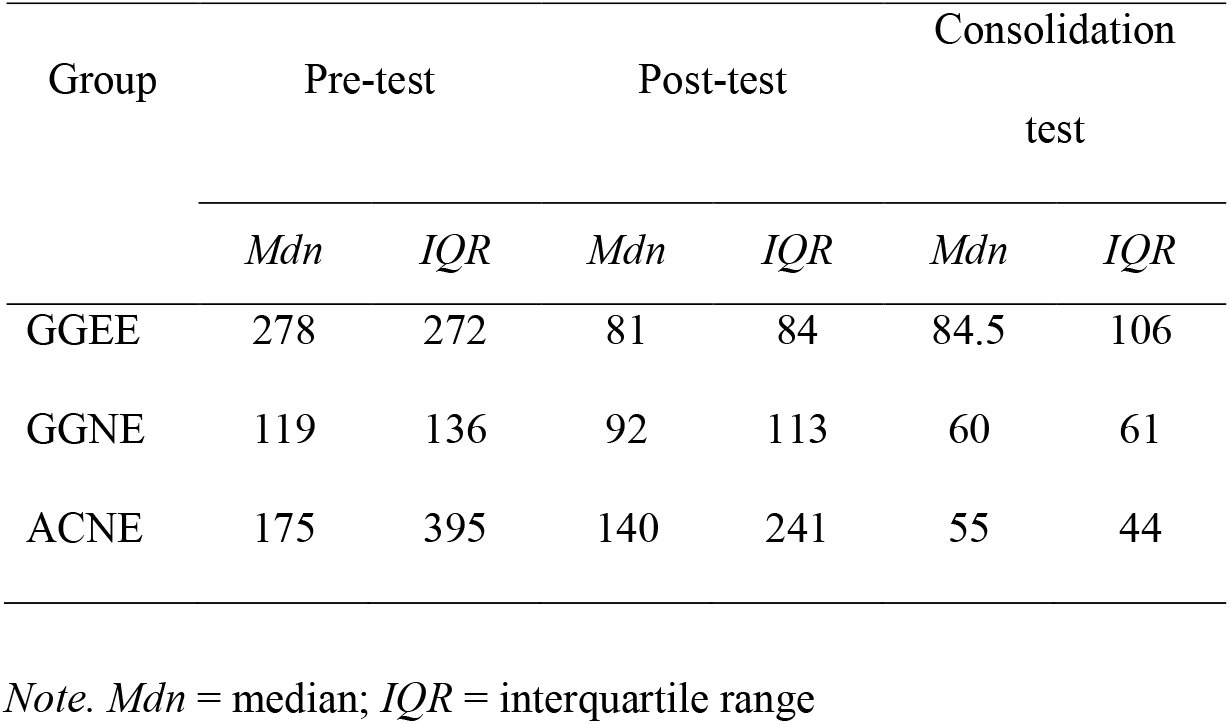
Descriptives of RTD Thresholds (in ms) for the Three Intervention Groups in the Three Separate Test Phases.

| Group | Pre-test |  | Post-test |  | Consolidation<br>test |  |
| --- | --- | --- | --- | --- | --- | --- |
|  | *Mdn* | *IQR* | *Mdn* | *IQR* | *Mdn* | *IQR* |
| GGEE | 278 | 272 | 81 | 84 | 84.5 | 106 |
| GGNE | 119 | 136 | 92 | 113 | 60 | 61 |
| ACNE | 175 | 395 | 140 | 241 | 55 | 44 |

*Note.* *Mdn* = median; *IQR* = interquartile range

**Figure 2.**
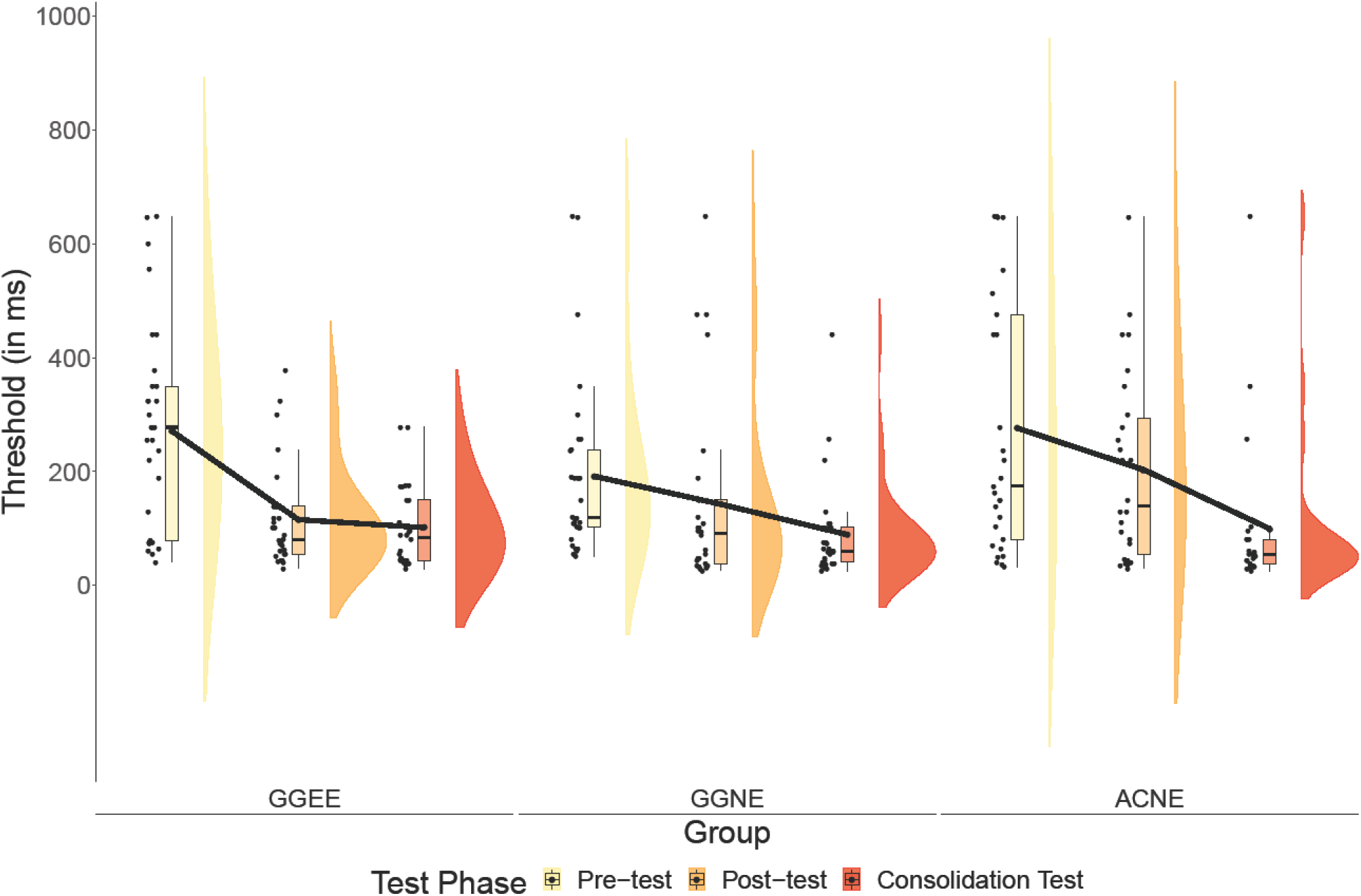
RTD Thresholds as a Function of Intervention Group and Test Phase. *Note*. From left to right we depict individual subjects’ thresholds, boxplots and probability distributions for the three intervention groups and the three test phases. The solid black line connects the means of the three intervention groups at each test phase.

**Figure 3.**
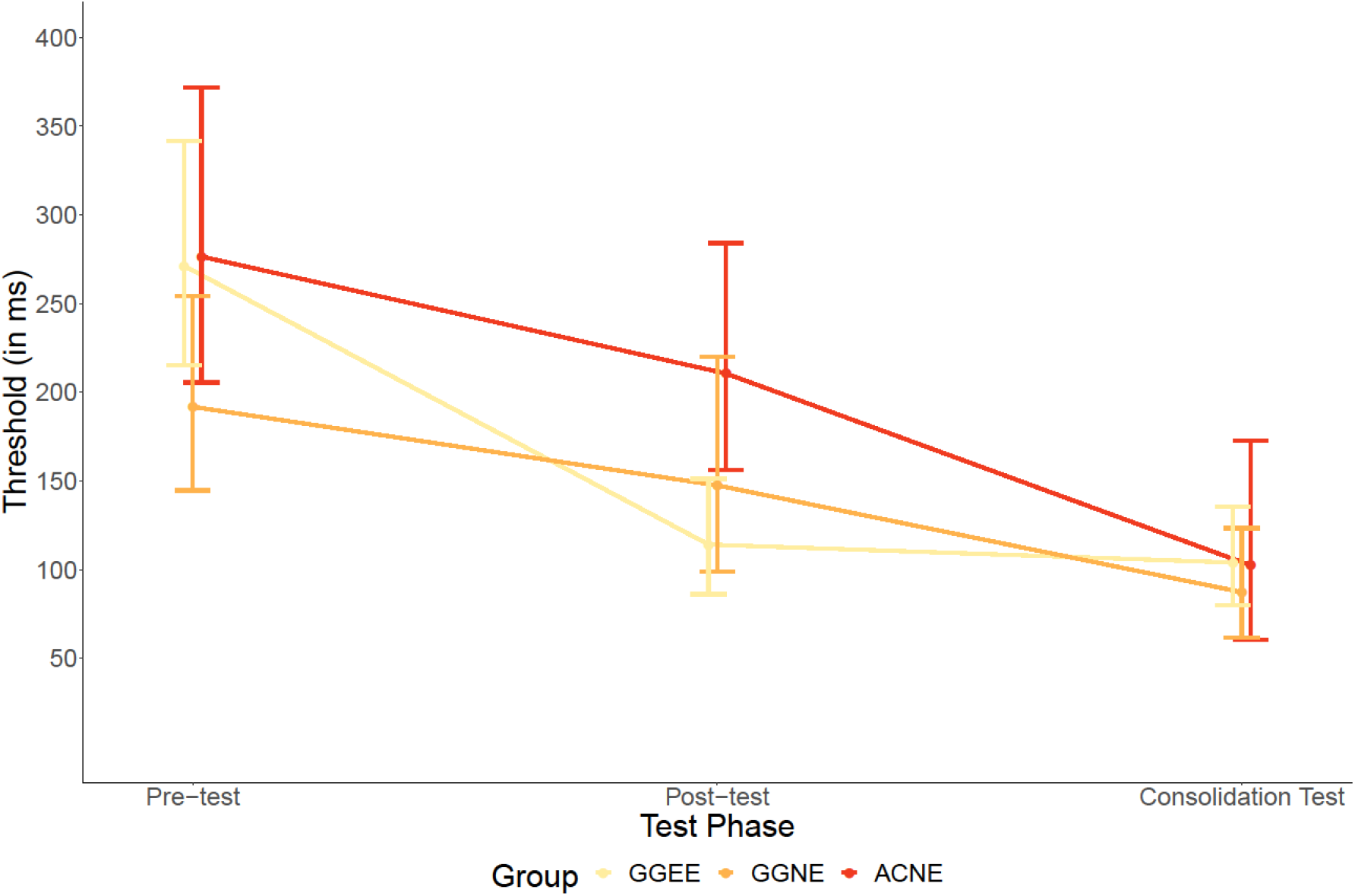
GEE Model Predictions. *Note*. Model predictions from the GEE model. Fitted values and 95% CIs are depicted for the three groups in the three test phases.

We furthermore compared the amount of hours the children in the different groups spent listening to the EE (GGEE group) or NE (GGNE and ACNE group) stories and the amount of hours they played GG (GGEE and GGNE group) or CG (ACNE group). The groups did not differ with respect to story listening (*F*(2,84) = 0.99, *p* = 0.37) or GG/CG (*F*(2.88) = 0.41, *p* = 0.67). See Table 3 for intervention duration descriptives.

**Table 3.** Descriptives of Hours of EE/NE Stories Listened to and Duration of GraphoGame and Control Game for the Three Intervention Groups.

| Intervention | GGEE |  | GGNE |  | ACNE |  |
| --- | --- | --- | --- | --- | --- | --- |
|  | *M* | *SD* | *M* | *SD* | *M* | *SD* |
| EE/NE Stories | 10.5 | 3.0 | 11.2 | 2.8 | 11.4 | 1.9 |
| GG/CG | 16.1 | 5.3 | 16.0 | 5.3 | 17.1 | 4.8 |

*Note.* *M* = mean; *SD* = standard deviation

## Discussion

The ‘dyslexia paradox’ often undermines the effectiveness of dyslexia remediation (Ozernov-Palchik & Gaab, 2016). In order to address this paradox, interventions should be administered preventively. Dyslexia remediation generally focuses on grapheme-phoneme correspondences with phonics-based interventions (National Institute of Child Health and Human Development, 2000). However, auditory processing difficulties in some children with dyslexia possibly supersede the beneficial impact of these interventions. This creates the need for interventions that (1) also target the auditory perceptive difficulties seen in dyslexia on top of the phonics-based interventions and that (2) can easily be administered preventively. Therefore, the present longitudinal study investigated the impact of a preventive combined auditory and phonics-based intervention in Dutch children at cognitive risk for dyslexia on an auditory temporal perceptive task, namely rise time discrimination. To our knowledge, this study is the first to investigate EE as an intervention strategy in prereaders at cognitive risk for dyslexia. Here, we demonstrated that the EE intervention in kindergarten directly boosts rise time discrimination, leading to improved rise time sensitivity at the start of reading acquisition while children without EE training only display rise time sensitivity improvement at a later moment in development.

We hypothesized that receiving the EE training would enhance children’s rise time sensitivity, whereas GraphoGame was not hypothesized to impact rise time discrimination. This EE-related improvement was expected to be demonstrated directly after the intervention, with potential long-term maintenance. Our results displayed an interesting distinction between the groups regarding the time period in which they mainly improved. In this study, EE provided an effect on rise time discrimination that could be measured immediately following the intervention, as demonstrated by the significant improvement following the intervention in the GGEE group. Previous studies by Van Hirtum, Moncada-Torres, et al. (2019) and Van Hirtum et al. (in press) already revealed the ability of EE to improve speech perception instantaneously and passively. The authors suggested that after long-term repeated exposure during intervention, EE might have an even stronger impact. In the current study we investigated the effect of such a repeated EE exposure on rise time discrimination and confirmed its impact on this measure. In contrast, the GGNE and ACNE groups did not seem to improve during the intervention period. Despite these marked differences in response to the intervention, the groups were not significantly different at post-test, although the difference between the GGEE and ACNE group tended towards significance, with the GGEE group outperforming the ACNE group. Even though we might have expected a clearer distinction between the group receiving the EE intervention and the groups not receiving this intervention, EE nonetheless provided a marked improvement that was not demonstrated by the other two groups. During first grade, when no intervention was taking place anymore, there was an improvement in the GGNE and ACNE groups only, hence they caught up with the GGEE group, reflected by no group differences at consolidation test. This decrease in the GGNE and ACNE groups seems to reflect maturation. Indeed, rise time discrimination has previously been found to improve with age in children with dyslexia, although to a lesser extent compared to typically developing children (Goswami et al., 2020). Apart from maturation, one might suggest that learning to read and the accompanying phonological awareness development improve auditory processing by refining acoustic contrasts in language (Kuppen et al., 2011). However, previous longitudinal studies have demonstrated that basic auditory processing impacts phonological awareness, but not the other way around, thereby contradicting this latter hypothesis (Goswami et al., 2020; Thomson et al., 2013). Our findings with respect to the phonics-based intervention further corroborate this statement and are in line with our expectations for the GraphoGame intervention. Considering the marked difference between the GGEE and GGNE groups, and the similar performance of the GGNE and ACNE groups, our phonics-based intervention did not seem to improve rise time discrimination. Hence, learning to read during the intervention did not impact rise time discrimination, making it unlikely that learning to read during first grade will exert an impact. This supports the maturation hypothesis of rise time discrimination. Considering that maturation in rise time discrimination performance has already been demonstrated until at least the age of 12 with a similar task to the one we used (Goswami et al., 2020; Kuppen & Goswami, 2016), we expected the GGEE group to also further improve during first grade hence maintaining its enhanced rise time sensitivity. The lack of further improvement in the GGEE group might be suggested to represent a floor effect that cannot be exceeded at this young age but might allow children to further improve at an older age. A fourth follow-up measurement would however be needed to confirm this hypothesis. However, the importance of our findings lies in the fact the EE intervention allows rise time discrimination to improve at a crucial time in development, namely right before the start of formal reading instruction. An issue with reading instruction, and as we previously suggested also with phonics-based interventions, is that they both rely on the accurate perception of presented phonemes. Hence, considering the auditory processing deficit, a necessary condition for learning to read, namely accurate phoneme perception, is not satisfied in some children with (a risk for) dyslexia. This could complicate their reading progress (Ziegler et al., 2009). The currently investigated EE intervention provided an improvement immediately following the intervention in the auditory processing of rise times, which provide important cues for phonetic discrimination (Goswami et al., 2011; Thomson et al., 2013). Accordingly, our intervention presumably has the potential to strengthen phoneme representations, which are of great importance for learning to read (Snowling & Hulme, 1994). Even more important, we demonstrated that the EE intervention allows for preventive intervention by implementation in children’s stories embedded in a game. More specifically, it shows potential to be implemented before the onset of formal reading instruction and thus during the most sensitive period for intervention (Ozernov-Palchik & Gaab, 2016). This enables us to intervene during a time period of great importance for brain plasticity (Gilmore et al., 2018), and a period in which the foundation is being set for later phonological awareness and reading development (Carroll et al., 2003; Snowling & Hulme, 2011). In this way, children receiving the EE intervention might have an advantage on the children not receiving this intervention at the onset of formal reading instruction. The latter group will start reading acquisition with unimproved rise time sensitivity that is possibly worse than that of typically developing children, which might complicate their reading progress, while for the children receiving the EE intervention this deficit is reduced by the time reading acquisition starts. Our EE intervention hence may tackle both the requirements we set forth for interventions, (1) it improves the auditory perceptive difficulties and (2) it can be administered preventively as to improve rise time discrimination at a crucial time in development.

Altogether, all groups clearly improved in rise time discrimination performance from pre-test (kindergarten) to consolidation test (end of first grade). We postulate that the improvement from kindergarten to the end of first grade in the GGNE and ACNE groups was the consequence of maturation during first grade, whereas in the GGEE group, the EE intervention provided a clear improvement, immediately following the intervention, which, importantly, even preceded the maturational effect in the other groups who only demonstrated this improvement during first grade.

### Limitations

Despite our promising findings the current study has a few limitations that need to be addressed. First, an important caveat is that it remains to be investigated whether the EE-induced improvement in rise time discrimination will in turn impact speech perception and phonological awareness in order to eventually improve reading performance in the investigated at risk children. However, previous longitudinal studies that have shown clear links between basic auditory processing such as rise time discrimination and later phonological awareness and reading allow us to hypothesize that this cascading effect should indeed be uncovered (Goswami et al., 2020; Vanvooren et al., 2017).

Another consideration is that in the current investigation there was no control group of typically developing children, and thus all children in this study are at cognitive risk for dyslexia. Consequently, we are unable to compare the developmental trajectories of typically developing children and children at cognitive risk for dyslexia. Previous longitudinal studies have already indicated differences between children with dyslexia and their controls in both the speed of rise time discrimination development (Goswami et al., 2020) and the trajectory of their development (Kuppen & Goswami, 2016). Notwithstanding the fact that our data are inadequate to directly test a possible differentiation between children at cognitive risk for dyslexia and typically developing children, the main message of our study remains that a preventive EE intervention allows for improved rise time discrimination performance immediately following the intervention. It might be suggested that EE will have a similar effect in typically developing children, considering its potential to also increase speech perception in children without dyslexia (Van Hirtum et al., in press), but this remains to be tested in future studies. However, administering the intervention to both typically developing and at risk children would not be recommended because it would possibly only enlarge the gap in performance between these groups.

## Conclusion

Our results show that speech envelope enhancement enables a direct improvement in basic auditory processing, in contrast to the maturation of these processes, which takes place at a later moment in development. Envelope enhancement affects basic auditory processing even before the onset of formal reading instruction. The significance here is that it allows for preventive interventions during the most effective intervention period. Further studies are required to investigate whether the improvement in rise time discrimination will also carry through to speech perception, phonological awareness and reading performance in order to mitigate (the occurrence of) dyslexia. Nevertheless, these results provide a first glimpse on a way to preventively tackle the auditory processing deficits that might be experienced by children at cognitive risk for dyslexia. Within a few years, we hope to have accumulated follow-up data from the children in this study. This will enable us to clearly distinguish between children that will have developed dyslexia as well as typically developing children. This will allow us to reanalyze our data by redefining the groups into dyslexic and typical reading groups and will even further clarify the significance of our EE intervention.

## Acknowledgements

We are most grateful to all children and parents for their cooperation in this study. Our special thanks goes to Ward Dehairs for the development of the EE/NE game, the speech-language therapists who recorded the stories for the EE/NE game and the stimuli for GraphoGame, colleagues at the Research Group ExpORL and Master students Speech Therapy and Audiology, Medicine and Educational Sciences that assisted in data acquisition.

## Ethics approval statement

This study was approved by the ethics committee of KU Leuven and written parental consent for participation of the children as well as oral assent by the children was obtained in line with the Declaration of Helsinki.

